# Photostimulation of lymphatic clearance of red blood cells from the mouse brain after intraventricular hemorrhage

**DOI:** 10.1101/2020.11.16.384149

**Authors:** Dong-Yu Li, Shao-Jun Liu, Ting-Ting Yu, Zhang Liu, Si-Lin Sun, Denis Bragin, Nikita Navolokin, Jürgen Kurths, Oxana Glushkovskaya-Semyachkina, Dan Zhu

## Abstract

Intraventricular hemorrhage (IVH) is the most fatal form of brain injury, yet a therapy directed at ameliorating intraventricular clot is very limited. There is accumulating evidence that an augmentation of the meningeal lymphatic (MLVs) functions might be a promising therapeutic target for IVH. In particular, the photostimulation (PS) of MLVs could be promising for non-invasive therapy of IVH via PS of clearance of red blood cells (RBCs) from the brain via MLVs. Indeed, we uncover that PS has therapeutic effects on IVH in mice reducing the mortality, improving the emotional status, accelerating the RBCs evacuation from the ventricles and increasing the ICP recovery. Our findings strongly suggest that the PS-mediated stimulation of drainage and clearing functions of MLVs can be a novel bedside, readily applicable and commercially viable technologies for treatment of IVH. These pilot results open new horizons in a non-invasive therapy of IVH via PS stimulation of regenerative lymphatic mechanisms.

## Introduction

Brain hemorrhage has the highest morbidity and mortality worldwide [1]. Intraventricular hemorrhage (IVH) is a bleeding into the brain’s ventricular system, where the cerebrospinal fluid (CSF) is produced and the circulates through towards the subarachnoid space. About 30% of IVHs are primary resulting from trauma and 70% are secondary, which originate from spontaneous intracranial hemorrage (ICH) in 45% and from aneurysmal of subarchnoid hemorrhage (SAH) in 25% [2–4]. Patients with IVH have a mortality from 50% to 80% and are twice as likely to have poor outcomes [5–7]. Blood in the ventricular system contributes to morbidity in a variety of ways. Pressure from the leaked blood damages brain cells make the damaged area unable to function properly. Besides, red blood cell (RBC) lysis after IVH results in the release of blood breakdown products (hemoglobin, iron, and bilirubin). Such products have been implicated in the pathophysiology of posthemorrhagic hydrocephalus and an increase in intracerebral pressure (ICP) because of impairment of CSF circulation and drainage system of the brain [8, 9]. Therefore, the conventional therapy of IVH, including surgery and fibrinolysis in combination with extraventricular drainage, centers on the blood evacuation from the ventricles, both for reducing ICP induced by mass effects of blood clots on the ventricular walls and the secondary damage caused by blood cell lysis [4, 10, 11]. However, the existing therapy of IVH has not made a dramatic impact on the natural history of the disease and new strategies are needed to reduce hematoma expansion and improving the drainage system of the brain [4, 10, 11].

One century ago, the hypothesis was proposed that the lymphatic pathway of clearance of red blood cells (RBCs) from the brain after ICH can be an important therapeutic target of brain hemorrhages [12–17]. In 1952, Simmonds demonstrated that RBCs can be cleared from the brain by lymphatic efflux [12]. The possibility of a lymphatic efflux of intracerebrally applied radioactively labeled RBCs has been described by authors elsewhere [13]. Oehmichen et al. found that intracerebrally applied RBCs move with CSF along connective pathway the subarachnoid space and accumulated in the cervical lymph nodes (cLNs) 1h after injection [14]. Caversaccio et al. [15] showed the intense detection of iron in the deep cervical lymph nodes (dcLNs) from patients suffering from ICH. Dontenwill described hemosiderin and fat phagocytosis in cLNs after a postnatal IVH [16]. Many studies using intracerebral injection of labeled blood cells (lymphocytes, macrophages, and RCBs) [13, 17], sarcoma cells [18], corpuscular tracers [19–22], water-soluble tracers such as ferritin and water-insoluble tracers such as lipoids [23, 24] demonstrated that the tracers reached cLNs along connecting fluid spaces, such as the perivascular space (PVS), or Virchow-Robin space (VRS) and the subarachnoid space, confirming the early experiments carried out by Simmond [12].

Quite recently the old concept of lymphatic pathway of RBC move from the brain was supplemented by new knowledge of RBCs clearance via the meningeal lymphatic vessels (MLVs) after ICH that was first proposed by our group in 2019 [25] and afterward it was studied after SAH by Chinese group in 2020 [26]. However, it still remain unanswered, whether RBCs can move from the ventricles into the lymphatic system and if so, whether this process can be therapeutically stimulated.

Photostimulation (PS), also known as low-level laser therapy (LLLT), can be a promising technology targeted to stimulation of brain drainage system for therapy of IVH. The transcranial PS is considered as a possible novel nonpharmacological and non-invasive therapy of stroke [27–29] and traumatic brain injuries [29–31]. The PS-mediated stimulation of the lymphatic drainage and clearing function might be one mechanisms playing an important role of PS in neurorehabilitation [32]. It has been proved that PS can regulate the relaxation and permeability of the mesenteric lymphatic vessels, activate moving of immune cells in the lymph, and has effectiveness in the management of lymphedema [33–35]. In our preliminary work, we clearly demonstrated that the near-infrared PS (1267 nm) stimulates clearance of tracers from the brain via MLVs after opening of the blood-brain barrier (BBB) or after its intracerebral injection [33, 34, 36–38]. We also showed that PS (1267 nm) effectively stimulates lymphatic clearance of beta-amyloid from the brain [39].

Based on these facts, we hypothesized that PS can effectively stimulate RBCs elimination from the ventricles and reduce pathological consequences after IVH. In this work, we studied the lymphatic pathway of RBCs clearance from brain after IVH and investigate whether PS (1267 nm) can enhance RBCs evacuation from the ventricles to improve the outcome after IVH. Using immunohistochemical and confocal colocalization analysis of the human and mouse brain samples, we clearly demonstrate the lymphatic clearance of RBCs from the brain in the post-hemorrhagic period. We also uncover that PS has therapeutic effects on IVH in mice reducing the mortality in 1.57 times, improving the emotional status, accelerating the RBCs evacuation from the ventricles and increasing the ICP recovery. Using coherence tomography and fluorescent microscopy for real time monitoring of PS stimulation of lymphatic clearance of tested tracers from the brain, we propose that PS stimulates drainage and clearing functions of MLVs that can be an important mechanism of PS-mediated managing of ICP and RCBs evacuation from the ventricles. Our findings opens new horizons in non-invasive therapy of IVH via PS stimulation of regenerative lymphatic mechanisms. The PS-mediated stimulation of drainage and clearing functions of MLVs can be a novel bedside, readily applicable and commercially viable technologies for effective routine treatment of IVH.

## Results

### The lymphatic clearance of RBCs from the human brain

In the first step, the presence of BRCs in MLVs and LVs of dcLNs taken from patients who died after IVH was studied. Our histological results reveled RBCs in both enlarged MLVs (87.21±6.10 μm vs. 39.33±1.12 μm, p<0.001, n=7) and in LVs of dcLNs (2713.07±21.18 μm vs. 1370.25±18.41 μm, p<0.01, n=7) in the next day after death due to IVH (Figure 1b, 1d and 1f). There were no RBCs in MLVs and in LV of dcLNs in the normal state (Figure 1a, 1c and 1d). Furthermore, the hemosiderin also was observed in dcLNs of patients after IVH, suggesting the RBC degradation in dcLNs that was not found in the normal lymph tissues (Fig. 1g and 1h).

**Figure 1.**
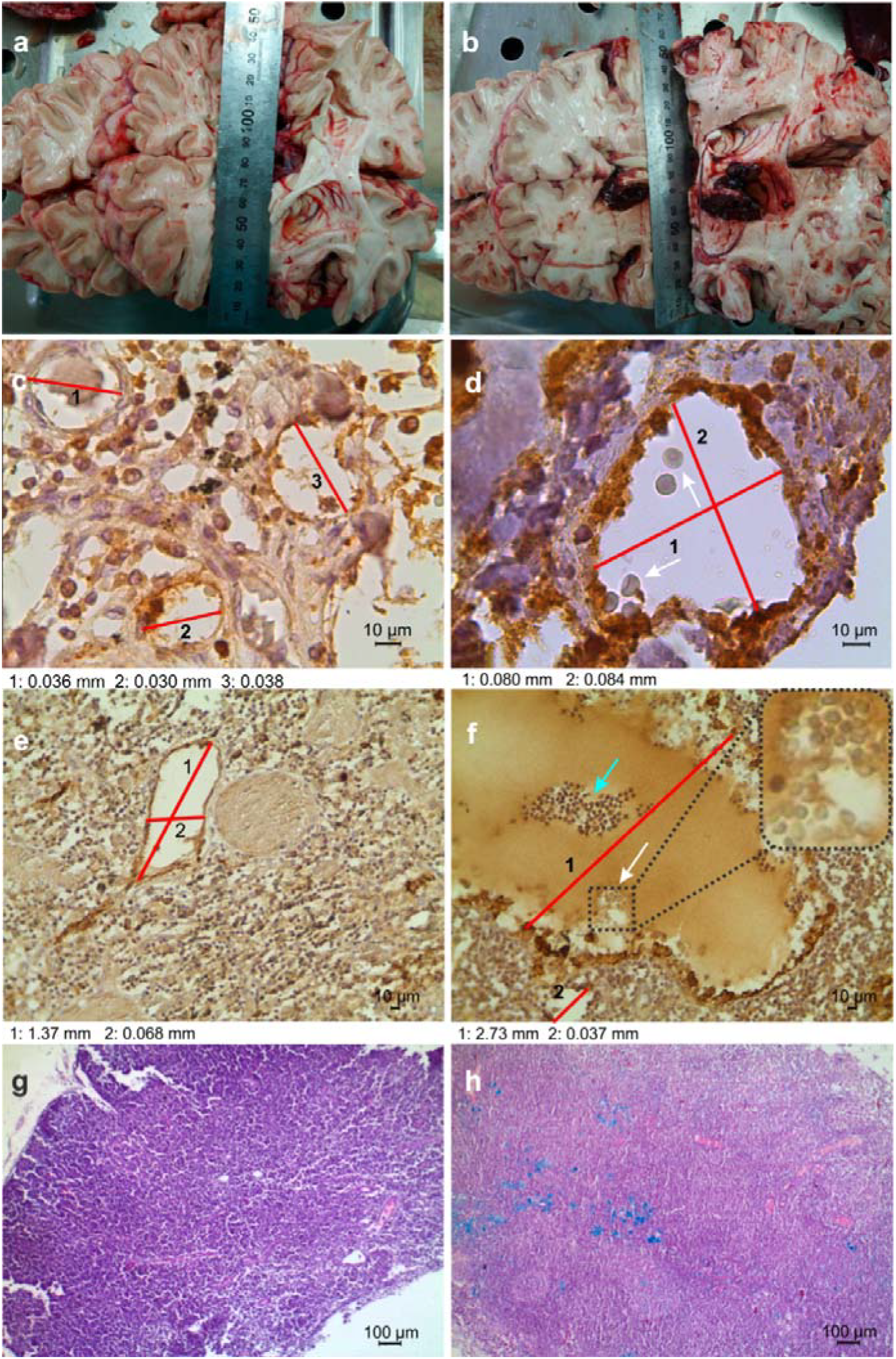
The lymphatic clearance of RBCs from the human brain after IVH: **(a)** and (**b)** - the images of the human brain without (a) and after IVH (b); **(c)** and **(d)** - Lyve-positive-labeled LVs (brown) in the dura matter without (c) and after IVH (d); **(e)** and **(f)** - Lyve-positive-labeled LVs in the dcLNs without (e) and after (f) IVH; **(g)** and **(h)** - the tissues of dcLNs without (g) and after IVH (h) (Hematoxylin □ Eosin). The bars: c and d - 774.0×; e and f - 246.4×, g and h - 64.6×The free RBCs (arrowed) and hemosiderin (blue color) are presented in the examined tissues taken in patients with IVH. The red lines in (с-f) show the diameters of LVs, n=7 in each group.

### The lymphatic clearance of RBCs from the mouse brain

Immunofluorescence and confocal colocalization analysis of markers of the lymphatic endothelium (Lyve-1 and Prox-1) and the blood vessels (CD-31) was used to study the meninges of mice without and after IVH (Fig. S1). Fig. 2a illustrates the absence of RBCs in MLVs in intact mice. The scatter plot in two directions, the Pearson’s correlation coefficient (PCC) value less than 0, and the small Mander’s value all point to that Red fluorescence is negatively correlated with green fluorescence. However, Fig. 2b clearly demonstrates the presence of RBCs in both MLVs and directly in the subarachnoid space, where arrows point out a series of RBCs entering LVs from the subarachnoid space. In addition, the Mander’s colocalization coefficients suggest that the Prox-1 colocalized well with Lyve-1, and nearly half number of RBCs were colocalized with LVs.

**Figure 2.**
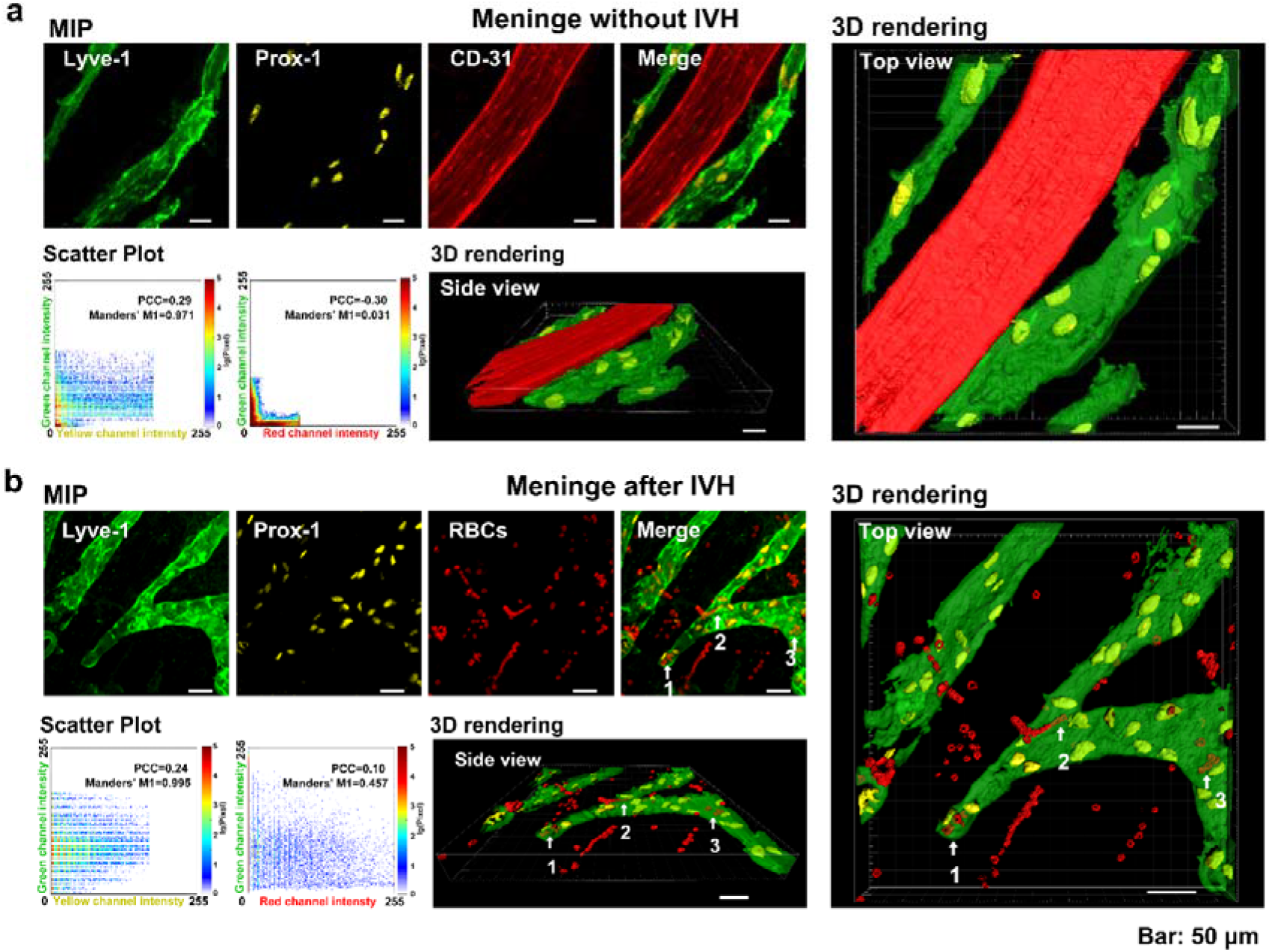
The clearance of RBCs from the mouse brain via MLS: Representative images of MLVs labelling by Lyve-1 (green color) and Prox-1 (yellow color) as well as the blood vessels labelling by CD31 (red color) without **(a)** and with IVH **(b)**. The 3D rendering images clearly shows that RBCs (red color, auto fluorescence) are either around MLVs or inside MLVs. MIP: Maximum intensity projection. PCC: Pearson’s correlation coefficient, which is between 1 and −1.1 represents perfect correlation, −1 represents completely negative correlation, and zero represents a random relationship. Manders’ M1: The proportion of the red/yellow fluorescence regions co-located with green

Furthermore, dcLN sections of mice with IVH was also visualized using confocal microscopy. As shown in Fig. 3a, red fluoresence signal could be hardly observed in LVs without IVH, which is consistent with the results of colocation analysis (PCC < 0, Mander’s colocalization coefficient of red signals v.s. green signals was as small as 0.01). However, as shown in Fig. 3b, RBCs were indeed observed inside LVs of dcLNs 3 hours after IVH, and nearly no RBC was outside LVs (Mander’s colocalization coefficient is 0.995).

**Figure 3.**
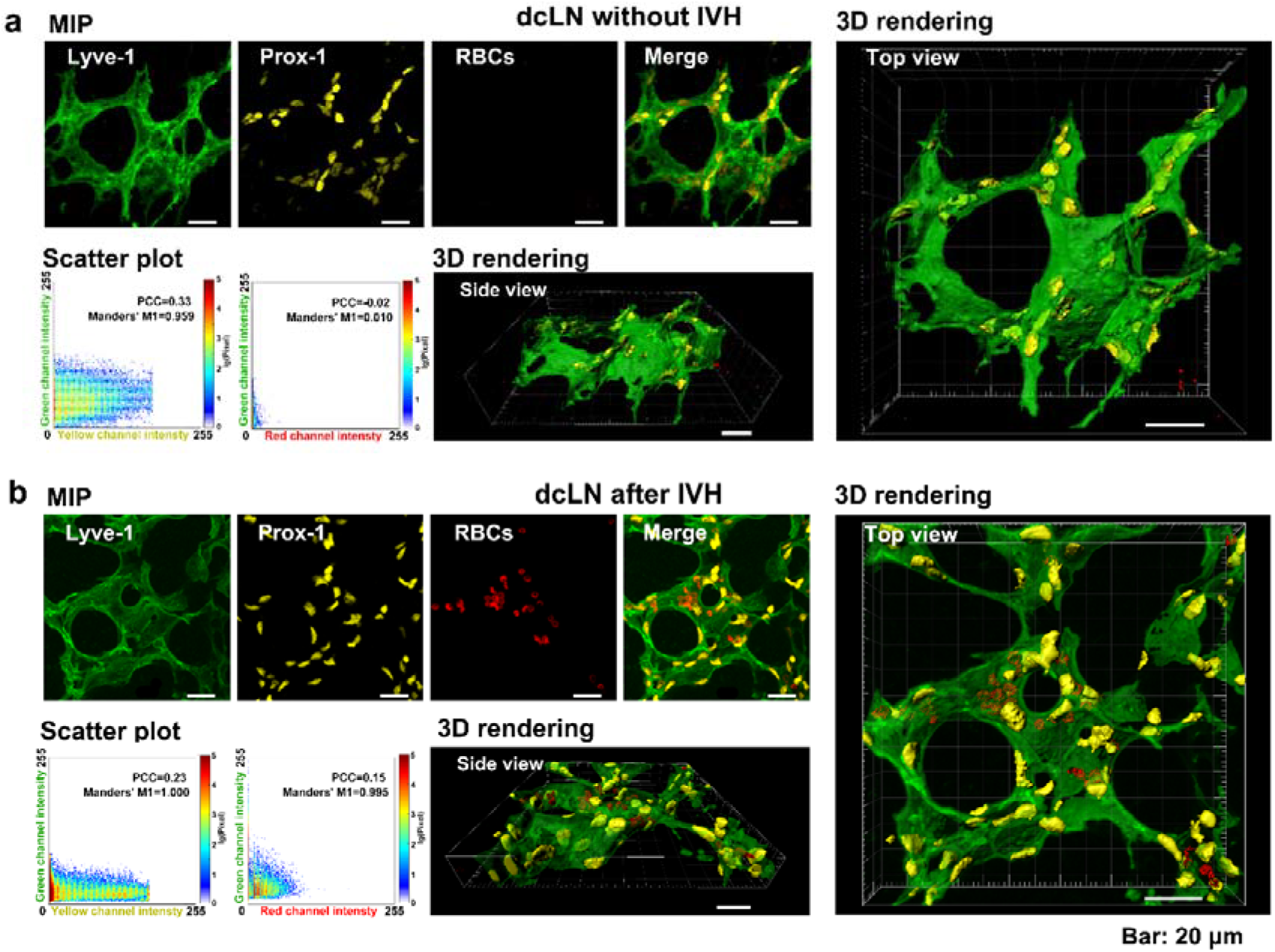
The clearance of RBCs from the mouse brain into dcLNs: Representative images of LVs of dcLN labelling by Lyve-1 (green color) and Prox-1 (yellow color) without **(a)** and after IVH **(b)**. The 3D rendering images shows RBCs inside LVs of dcLN. MIP: Maximum intensity projection.

Thus, both human and animal data clearly suggest the lymphatic pathway of RBCs clearance from the brain into dcLNs, which is consistent with the oldest and latest data about lymphatic efflux of RBCs [12, 17, 25, 26].

### PS increases the diameter and the distribution of MLVs

As reported in our previous studies of PS-mediated stimulation of meningeal lymphatic clearance and drainage function, the 1267-nm laser in a dose of 9-10 J/cm^2^ (17 min of laser irradiation, 5-min pause, 61 min in total) is effective for lymphatic clearance of small and large molecules from the brain into dcLNs [33, 34, 39]. Based on our pervious animal data and the established PS protocol [33, 34, 39], we investigated the effect of PS on MLVs using a 1267-nm laser with an indicated above dose. We uncovered that although IVH seemed to widen the diameter distribution of MLVs, it didn’t make significant difference on the average diameter (25.8±18.2 μm vs 25.75±15.7 μm). However, NIR PS enlarged the diameter of MLVs after IVH (25.75±15.7 μm vs 31.0±12.8 μm, p<0.01, Fig. 4). These *in vivo* results indirectly suggest that PS could enhance the permeability of MLVs via relaxation of the lymphatic endothelium that has been shown in our preliminary data [34].

**Figure 4.**
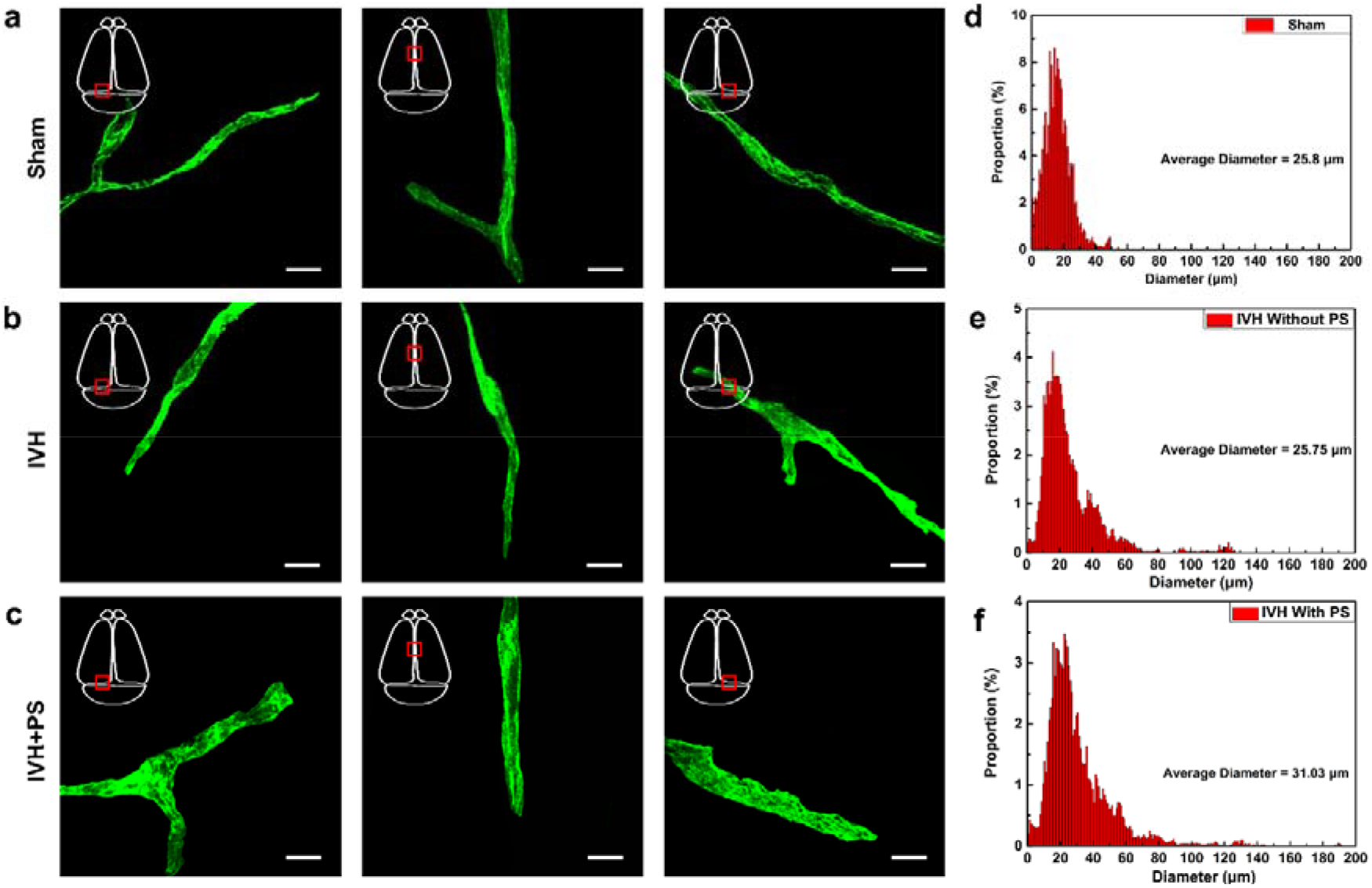
The PS-mediated enlargement of MLVs in healthy mice: **(a-c)** Representative fluorescent images of MLVs (labelled with lyve-1) in the transverse sinus of sham group (a), IHV group (b) and IHV+PS group (c); **(d-f)** The distribution of MLVs in sham group (d), IHV group (e) and IHV+PS group (f), n=10 in each group. Bar: 20 μm.

### PS stimulates lymphatic clearance of RBCs and tested tracers from the brain

In *in vivo* experiments on healthy mice, we study the effectiveness of the selected dose of PS for stimulation of lymphatic clearance of tracers (Evans Blue dye (EBD) and gold nanorods (GNRs)) from the right lateral ventricle mimicking the road of RBCs elimination from the brain. Using light tracers from the brain into dcLNs (Fig. 5 and S2). Indeed, the intensity of EBD fluorescence was 2.6-fold higher in mice treated by PS vs. intact animals (11.5±4.5 vs. 4.5±2.0, p<0.05) (Fig. 5a and 5b). Using optical coherence tomography (OCT), we found that the rate of GNRs accumulation in dcLNs is extremely small in the control group (no PS) and is 1.4-fold higher after PS (104±5.3 a.u. vs. 144±6.9 a.u. at 60 min of observation, p<0.001, Fig. 5c). The *ex vivo* results of atomic absorption spectroscopy confirmed the OCT data and showed that the level of GNRs in dcLNs is 3.1-fold higher in the PS-group vs. the control group (4.0±0.9 μg/g tissue vs. 12.7±1.7 μg/g tissue, p<0.001). Thus, our results on healthy mice confirmed our previous data suggesting that PS 9 J/cm^2^ (17 min of laser irradiation, 5-min pause, 61 min in total) effective stimulates lymphatic clearance of tracers injected into the right ventricles.

**Figure 5.**
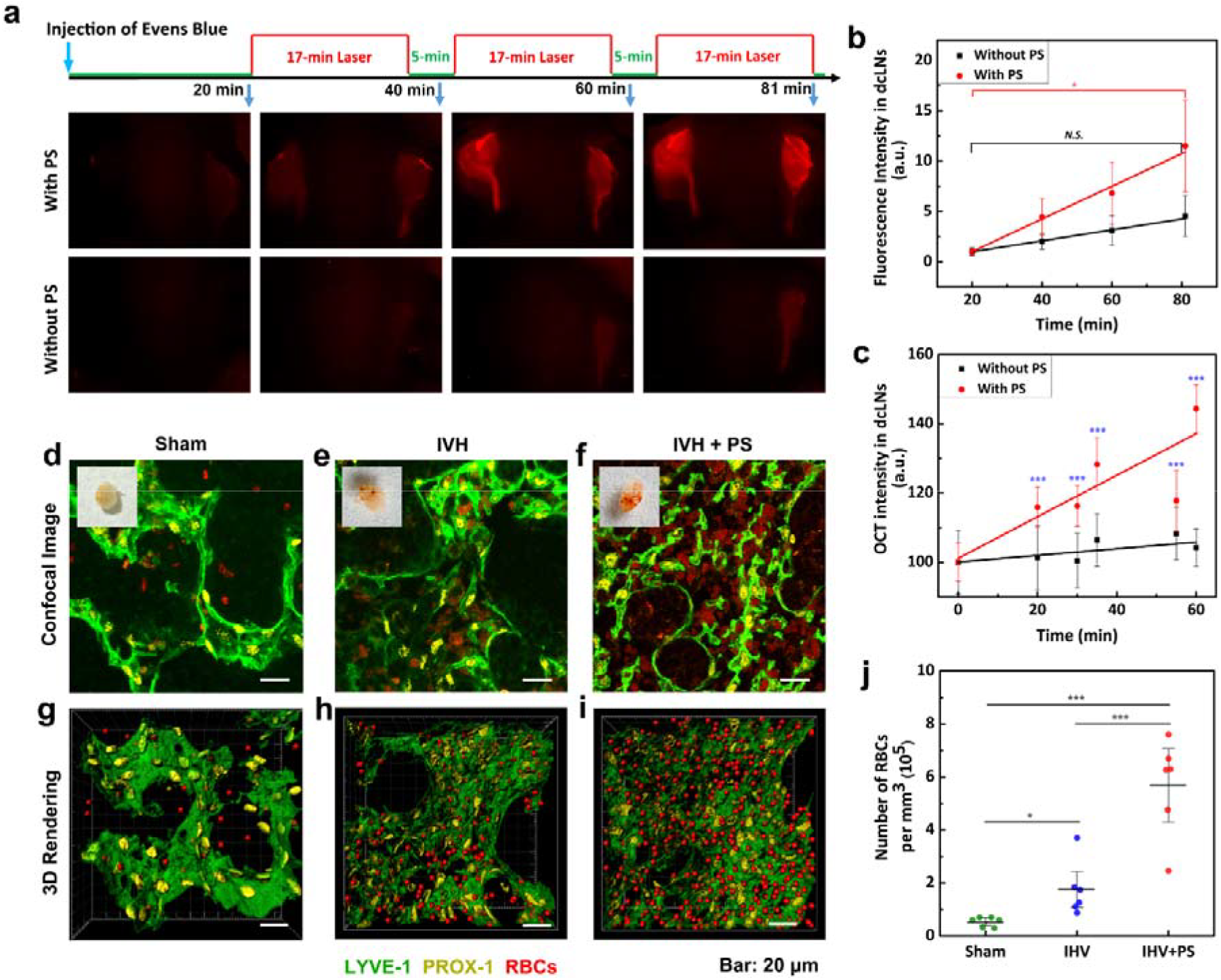
The PS stimulation of lymphatic clearance of tracers in intact mice and RBCs in mice after IVH: **(a**) Representative fluorescent images of EBD clearance from the right ventricle into dcLNs with and without PS; (**b**) Quantitative analysis of fluorescence intensity of EBD accumulation in dcLNs with and without PS; (**c**) the OCT data of GNRs accumulation in dcLNs after its intraventricular injection with and without PS, * - p<0.05, n=10 in each group of all series of experiments; **(d)** Typical confocal images of RBCs in LVs of dcLN 1 hour after saline injection; **(e)** Typical confocal images of RBCs in LVs of dcLN 1 hour after blood injection **(f)** Typical confocal images of RBCs in LVs of dcLN 1 hour after blood injection with laser irradiation. The insets in (d)-(e) represent macro photos of dcLN; **(g)** - **(i)** 3D rendering of distribution of RBCs in LVs of dcLN 1 hour after saline injection (g), IVH without PS (f) and IVH PS (g) (the volume of dcLN was 135×135×40 μm^3^); **(j)** the number of RBCs in dcLN 1 hour after IVH with and without NIR PS, * - p<0.05, *** - p<0.001,n=6 in each group. The LVs were labeled by Lyve-1 (green color) and Prox-1 (yellow color), the RBCs were imaged with its autofluorescence (red color).

Using the established protocol of PS on healthy mice, in the next step, we studied PS effects on RBCs clearance from the brain in mice after IVH. The results presented in Fig. 5d-5i clearly demonstrate that the number of RBCs in dcLNs is significantly higher after PS irradiation compared with the IVH group and sham group ((0.5±0.1)×10^5^ vs. (1.7±0.7)×10^5^ vs. (5.7±1.4)×10^5^ per mm^3^, p<0.001).

### PS therapy of IVH: the effect on the intracranial pressure (ICP), the mortality rate and emotional status

In final step, we studied possible therapeutic effects of PS after IVH in mice. Figure 6a clearly demonstrates that IVH is accompanied by a dramatical rise of ICP that was significantly improved by PS. So, the blood injection into the right ventricle immediately caused an increase in ICP (75.5 ± 12.5 mm Hg and 74.2 ± 11.4 mm Hg vs.10.2 ± 2.1 mm Hg, p<0.01 for the sham and the IVH/IVH+PS groups, respectively). Afterward, ICP gradually decreased but remained to be high by the end of 60 min monitoring. The PS significantly reduced the time of ICP recovery. Indeed, the recovery of ICP in the IVH+PS group was faster than in the IVH group (29.0 ± 3.1 vs. 19.6 ± 3.2 mm Hg at 60 min of monitoring, respectively, p<0.05).

**Figure 6.**
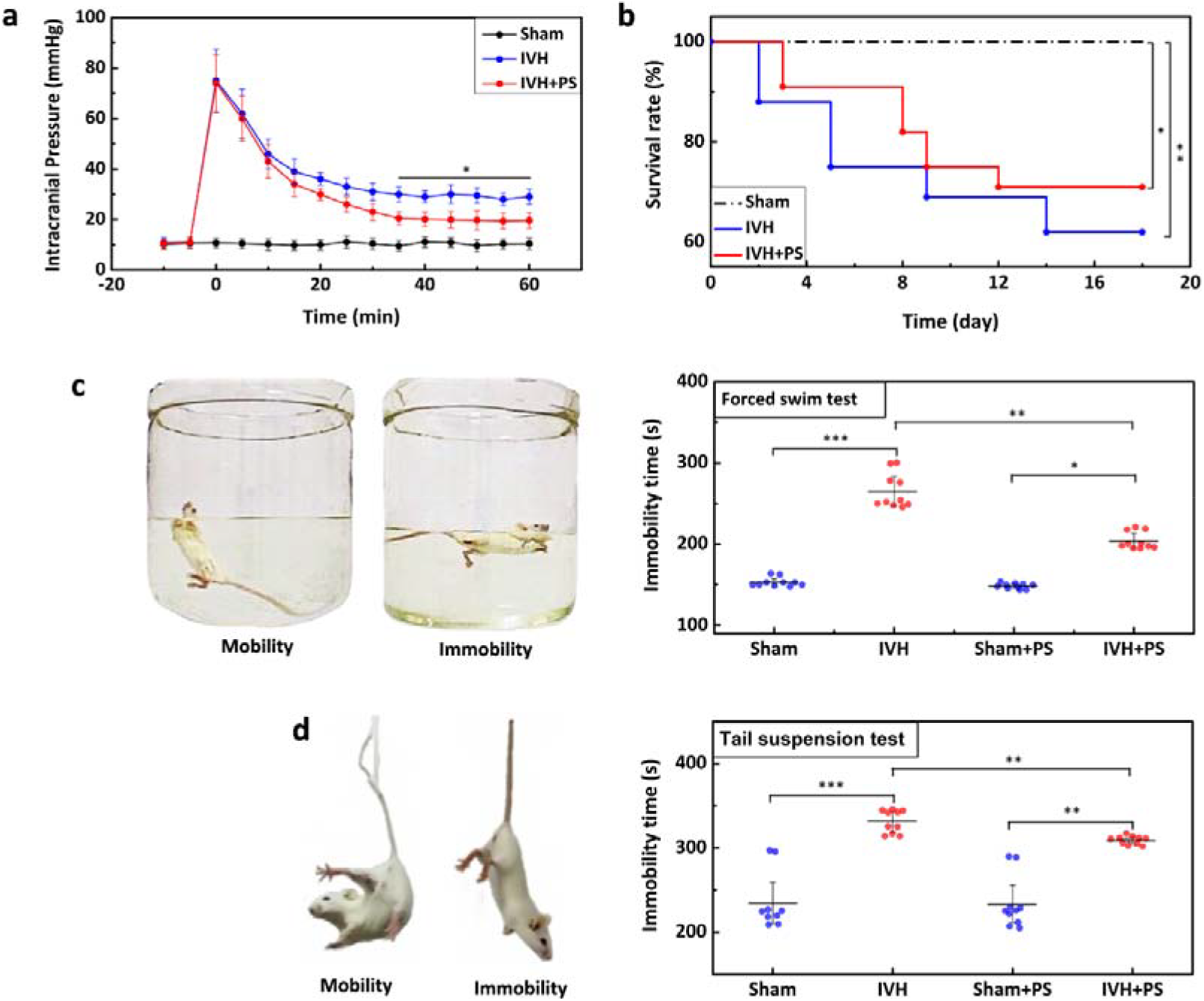
The PS effects on ICP, the mortality rate and emotional status in mice from the sham, IVH/IVH+PS groups: (**a**) **–**The continues monitoring of ICP after IVH with and without PS, * - p<0.05 vs. the baseline level; (**b**) – The mortality rate in mice after IVH with and without PS; **(c)** and **(d)** The immobility time in the tail suspension (c) and in the forced swim test (d); Mean ± SD; *** - p<0.001; ** - p<0.01; * - p<0.05, n=30 per group in (b) and n=10 per group in a, c and d.

Since the recovery after IVH requires a long-term therapy, we studied the effects of PS course (9 J/cm^2^ on the surface of the brain, 17 min of laser irradiation, 5 min - pause, during 61 min, during 7 days, each second day) on such vital signs as the mortality and emotional disorders in mice after IVH. The mortality of mice was 36.7% (11 of 30) in the IVH group. No animals died in the sham groups. The PS course increased the survival of mice. So, the mortality in the IVH+PS group was 23.3% (7 of 30), i.e. 1.57-fold lesser than in IVH group (Fig. 6b).

The tail suspension and forced swim tests were used for evaluating the emotional abnormality after IVH. The time of immobility in the tail suspension and forced swim tests was longer in the IVH than in the sham group (n=10 in each group, p<0.001; Fig. 6c and 6d). The PS course was accompanied by a significant decrease of immobility time in both tail suspension and forced swim tests compared to the IVH group (p<0.05, n=10 in each group). The PS did not affect the performance of both tests in the sham group.

Thus, these results suggest that PS contributes to a faster recovery of ICP after IVH and the PS-course causes 1.57-fold decreases of the mortality and improves the emotional status of mice with IVH.

## Discussion

In this work, we studied the lymphatic pathway of RBCs clearance from brain after IVH and investigated, whether PS (1267 nm) can enhance RBCs evacuation from the ventricles to improve the outcome after IVH. In the *ex vivo* experiments using human and mouse brain samples, we clearly demonstrated that RBCs move from the ventricles via MLVs into dcLNs. Thus, our findings confirm the lymphatic clearance of RBCs from the brain in post-hemorrhagic period. These results provide strong support for the old theory of RBCs evacuation from the brain via lymphatic efflux [12–17]. Our data are also consistent with the newest discovery of the meningeal lymphatic pathway of clearance of RCBs from the brain after brain hemorrhage [25, 26]. Taking into account all of the above, we hypothesized that the lymphatic mechanisms of blood clearance from the brain can be a new therapeutic target for treatment of IVH.

To test our hypothesis, we investigated whether PS (1267 nm) can enhance RBCs evacuation from the ventricles to improve the outcome after IVH. Here we chose a wavelength of 1267 nm, which belongs to NIR-II region and was not in common use for PS. For transcranial PS, the light should successively penetrate scale, skull and the brain tissue above the targeted layer. However, the tissues have severe absorption and scattering to light, which may limit the efficacy of PS. Because the tissue scattering decreases with the wavelength increases, NIR light, such as 808 nm, 980 nm, 1064 nm and 1072 nm [27–32], instead of visible light, has been more and more used for transcranial PS [40]. Compared to those wavelengths, ~1300 nm suffers even less scattering and causes less photodamage to tissues, which has been used for transcranial deep-tissue imaging [41]. Therefore, such wavelength is reasonable to be considered for transcranial PS, while relative work has seldom been reported.

Herein, Based on our preliminary animal data and the established PS protocol (the 1267-nm laser in a dose of 9-10 J/cm2, 17 min of laser irradiation, 5-min pause, 61 min in total) [33, 34, 39], we investigated the effect of PS on MLVs without and after IVH.

In *in vivo* experiments on mice, we uncover that PS has therapeutic effects on IVH in mice reducing the mortality in 1.57 times, improving the emotional status, accelerating the RBCs evacuation from the ventricles and increasing the ICP recovery. Our results are in agreements with other animal and human data suggesting a greater clinical significance of transcranial near-infrared laser phototherapy of stroke and brain trauma [27–31].

To study mechanisms of PS therapeutic effects, we used coherence tomography and fluorescent microscopy for real time monitoring of PS stimulation of lymphatic clearance of tested tracers from the ventricle. We clearly demonstrated that the accumulation of EBD and GNRs in dcLNs after its intraventrical injected was 2.6-fold, p<0.005 greater (EBD) and 1.4-fold higher, p<0.001 (GNRs) in the PS group vs. the control group (no PS). These findings allowed us to propose that PS stimulates drainage and clearing functions of MLVs and can be an important mechanism of PS-mediated managing of ICP and RCBs evacuation from the ventricles. Indeed, our confocal data demonstrate that PS enlarges MLVs and increases its distribution in the meninges suggesting PS-mediated influences on the tone of the lymphatic vessels. In our preliminary data we revealed that PS causes relaxation of the mesenteric lymphatic endothelium that is associated with an increase permeability of lymphatic walls and a decrease of expression of tight junction proteins [34]. Probably, these effects can be related to a PS-mediated increase in the activity of endothelial nitric oxide (NO) synthase [42]. NO is a vasodilator that acts via stimulation of soluble guanylate cyclase to form cyclic-GMP (cGMP), which activates protein kinase G causing the opening of calcium-activated potassium channels and re-uptake of Ca^2+^. The decrease in concentration of Ca^2+^ prevents myosin light-chain kinase from phosphorylating the myosin molecule, leading to relaxation of lymphatic vessels [43]. There are several other mechanisms by which NO could induce lymphatic dilation: 1) the activation of iron-regulatory factor in macrophages [44], 2) the modulation of proteins such as ribonucleotide reductase [45] and aconitase [46]; the stimulation of the ADP-ribosylation of glyceraldehyde-3-phosphate dehydrogenase [47] and protein-sulfhydryl-group nitrosylation [48]. We assume that the PS effects on the tone of MLVs facilitate drainage in fluid spaces of the brain that drains RBCs/tracers from the ventricles along the subarachnoid space, where RBCs partly penetrate into MLVs. We also do not exclude the idea of a possible presence of the putative lymphatic vessels directly in the brain tissues that has been presented by Prineas in 1979 on the human brain samples obtained from patients who died from various neurological disorders [49].

Our findings opens new horizons in a non-invasive therapy of IVH via PS stimulation of regenerative lymphatic mechanisms. The PS-mediated stimulation of drainage and clearing functions of MLVs can be a novel bedside, readily applicable and commercially viable technologies for effective routine treatment of IVH and other types of brain bleedings. Our pilot experiments were focused on demonstrating the phenomenon of RBCs clearance via MLVs and PS stimulation of this process that needs further detailed investigations of clinical efficiency of PS-mediated stimulation of lymphatic functions in the human brain. Due to scattering effects of the skull, the laser penetration into the brain is very limited that is most complication of PS. However, we strongly believe that PS-acceleration of RBCs clearance from the brain via MLVs can be clinically important for a therapy of brain hemorrhages in newborns, in which PS can be applied through a special window into the brain as the fontanelle. Note, that IVH occurs in 45% of premature infants [50, 51]. The PS-lymphatic therapy can be important also for silent SAH and subdural hemorrhages, which are located superficially and are identified in 46-50% of mature babies [52]. Our data are limited by the study of one protocol of PS influences on the lymphatic evacuation of RBCs, while it has actively discussed that sleep can be a natural factor of activation of the lymphatic functions in the brain [53]. We assume that PS application during deep sleep can significantly enhances therapeutic effects of PS on brain bleedings and causes more pronounced influences on regenerative lymphatic mechanisms.

## Methods

### Subjects

Autopsy specimens of human dura and the dcLNs were obtained from the Department of Pathological Anatomy at the Saratov State University (average age 42). All obtained samples were fixed and stored in a 10% formalin solution for prolonged time periods. The instigations were performed on the following groups: 1) the control - healthy patients died after fatal traumatic injuries (n=9); 2) patients died after IVH, n=15.

Male BALB/c mice (20–25 g) were used in all experiments. The animals were housed under standard laboratory conditions, with access to food and water, *ad libitum*. All procedures were performed in accordance with the “Guide for the Care and Use of Laboratory Animals”. The experimental protocols were approved by the Local Bioethics Commission of the Saratov State University (Protocol No. 7); Experimental Animal Management Ordinance of Hubei Province, P. R. China (No. 1000639903375); the Institutional Animal Care and Use Committee of the University of New Mexico, USA (#200247). The animal experiments included following groups: 1) the intact, healthy mice; 2) the IVH group were injected the autologous blood into the right lateral ventricle; 3) sham control mice were injected an equal volume of saline; 4) sham + photostimulation (PS) group, and 5) the IVH group+PS.

### Immunohistochemistry for the human samples

For visualization of human LVs in the dura matter and the dcLNs, the protocol for the immunohistochemical (IHC) analysis was used with the markers Lyve-1 (eBioscience, San Diego, USA). Briefly, tissue samples were fixed with formaldehyde and, after routine processing, were embedded into a paraffin block. Then the samples were sectioned into 3-to 5-μm slides; afterward, they were dried at 37°C for 24 h and then rehydrated by sequential incubation in xylene (three times, 3 min each), 96% ethanol (three times, 3 min each), 80% ethanol (two times, 3 min each), and distilled water (three times, 3 min each). The IHC reaction was visualized with a REVEAL— BiotinFree Polyvalent diaminobenzidine kit (Spring Bioscience). Endogenous peroxidases were blocked by adding 0.3% hydrogen peroxide to the sections for 10 min, followed by washing of sections in phosphate-buffered saline (PBS). The antigen retrieval was conducted using a microwave oven in an ethylenediaminetetraacetic acid-buffer pH 9.0, and a nonspecific background staining was blocked in PBS containing 0.5% bovine serum albumin and 0.5% casein for 10 min, after which the sections were washed in PBS for 5 min. Further, the sections were incubated in a humid chamber with diluted anti-Lyve-1 (eBioscience, San Diego, USA; 1L1000) for 1 h at room temperature. After that, the sections were washed in PBS, incubated with secondary horseradish peroxidase-labeled goat antirabbit antibodies for 15 min, again washed in PBS, counterstained with hematoxylin for 1 min, washed again in water, dehydrated in graded alcohols (three times, 3 min each) and then in xylene (three times, 3 min each), and finally embedded into Canadian balm.

Hemosiderin was identified histologically with Perls’ Prussian-blue stain, where iron in hemosiderin turns blue to black when exposed to potassium ferrocyanide. Since hemosiderin is an iron-storage complex (denatured ferritin and other material), which is marker of organ damage associated with hemorrhages [54, 55].

### Microscopy for human sections

All human sections were imaged by white light microscopy using the digital image analysis system Mikrovizor medical μVizo-103(LOMO, Russia).

### Modeling of intraventricular hemorrhage in mice

To produce the mice IVH model, autologous blood was injected into the right lateral ventricle. Aseptic techniques were used in all surgical procedures. The disinfection with Betadine and 70% ethanol of the stereotactic apparatus and surgical tools was made prior to surgery. Throughout surgery and the experimental period, rectal temperature was monitored until mice completely recovered and displayed normal motor activity. The ketamine (100 mg/kg) and xylazine (10 mg/kg) was injected intraperitoneally for the anesthesia. The mouse was placed onto a thermal blanket and the scalp was shaved. The ophthalmic ointment to both eyes was applied. A 1 cm long midline incision of the scalp with a 10 scalpel blade was made. The Hamilton syringe (25 μl) was mounted onto the injection pump, and the needle (25 Gauge) over bregma was directed stereotaxically. Next, the needle was positioned 0.5 mm posterior and 1.06 mm lateral of the bregma to the right with the stereotactic manipulator. A small cranial burr hole was drilled through the skull using a variable speed drill with a 1 mm drill bit. The animal’s tail was disinfected with 70% ethanol and the central tail artery was punctured with a sterile needle (25 Gauge), then the arterial blood was fixed with heparin sodium and collected into a capillary tube. The 10 μl of blood was quickly transferred from the capillary tube into the glass barrel of the Hamilton syringe and inserted the plunger. The needle was inserted into the right lateral ventricle to a depth of 2.5 mm below the skull surface. The arterial blood was injected at a rate of 2 μl/min. The needle was left in the ventricle for 10 min and then removed at a rate of 1 mm/min to prevent the reflux of blood. The burr hole and scalp incision were closed with bone wax (Ethicon, Somerville, NJ) and with cyanoacrylate glue (Henkel Consumer Adhesive Inc. Scottsdale, Arizona), respectively. Sham control mice were injected with an equal volume (10 μl) of saline.

### Photostimulation of lymphatic system of brain

The mice with shaved head were fixed in a stereotaxic frame and irradiated in the area of the parietal cortex in the region of the Sagittal sinus using the sequence of: 17 min – irradiation, 5 min – pause, 61 min in total. A laser diode (LD-1267-FBG-350, Innolume, Dortmund, Germany) emitting at 1267 nm was used as a source of irradiation. The laser diode was pigtailed with a single-mode distal fiber ended by the collimation optics to provide a 5 mm beam diameter at the specimen. The laser dose of 60 J/cm^2^ was used in this work. It was previously reported that 15% of the laser energy goes to the brain via the intact mouse skull [33]. Therefore, the power density at 9 J/cm^2^ was on the surface of the mice brain. The heating of the brain tissue caused by exposure to light was monitored by using a thermocouple data logger (Pico Technology, USB TC-08, Cambridge shire, UK).

To exclude thermal effects of PS on the brain, preliminary we analyzed the changes in temperature on the surface of the brain before and after PS. To measure the brain surface temperature, the medial part of the left temporal muscle was detached from the skull bone, a small burr hole was drilled into the temporal bone, and a flexible thermocouple probe (IT-23, 0.23 mm diam, Physitemp Instruments LLC, NJ, USA) was introduced between the parietal bone and brain into the epidural space. The brain surface temperature was measured before, and during the laser application using a handheld thermometer (BAT-7001H, Physitemp Instruments LLC, NJ, USA). There were no any changes in the temperature of the brain surface after PS (37.15±0.11 vs. 37.13±0.56).

### Low-power laser therapy after IVH

For immediate treatment, the mice were treated with photostimulation right after surgery procedure of blood injection into the right lateral ventricle, followed by the removal of dcLNs to measure the RBCs accumulation in the lymphatic system.

For long-term therapy, the mice were treated by PS during 7 days each second day under inhalation anesthesia (1% isoflurane at 1 L/min N_2_O⁄O_2_ - 70:30) 3 days after the surgery procedure of blood injection into the right lateral ventricle (Fig. S3), during which period the survival and behavioral assessment were performed.

### Immunohistochemistry for the mouse tissue

To visualize the lymphatic system and vasculature of mouse meninges and dcLN, fluorescent markers were used to label the certain structures using immunohistochemical method [56]. Anti-Lyve-1 and anti-Prox-1 were used to label lymphatic vessels; anti-CD-31 was used to label blood vessels.

Mice were sacrificed with cervical dislocation. To obtain the meninges, skin was removed from the head and the muscle was stripped from the bone. After removal of the mandibles and the skull rostral to maxillae, the top of the skull was removed with surgical scissors. Whole-mount meninges were fixed while still attached to the skull cap in PBS with 2% paraformaldehyde (PFA) over night at 4 □. The meninge was then dissected from the skullcap. For analysis of the dcLNs, the lymph nodes were removed and fixed in PBS with 2% PFA over night at 4 □, and then fixed in 2% agarose, followed by sliced into 60 μm-thick sections using a vibratome (Leica VT1000, Germany).

The whole mounts of meninges and the sections of dcLNs were firstly washed 3 times (5 min for each) with wash solution (0.2% Triton-X-100 in PBS), secondly incubated in the blocking solution (a mixture of 2% Triton-X-100 and 5% normal goat serum in PBS) for 1 hour, followed by incubation with Alexa Fluor 488-conjugated anti-Lyve-1 antibody (1:500; FAB2125G, R&D Systems, Minneapolis, Minnesota, USA), rabbit anti-Prox-1 antibody (1:500; ab 101851, Abcam, Cambridge, United Kingdom) and mouse Alexa Fluor 647-conjugated anti-CD31 antibody (1:500; 102416 BioLegend, San Diego, USA) overnight at 4°C in PBS containing 0.2% Triton-X-100 and 0.5% normal goat serum. Next, the meninges were incubated at room temperature for 1 hour and then washed 3 times, followed by incubation with goat anti-rabbit IgG (H+L) Alexa Flour 561 (Invitrogen, Molecular Probes, Eugene, Oregon, USA).

### Confocal microscopy

The mouse meninges and dcLN sections were imaged using a confocal microscope (LSM 710, Zeiss, Jena, Germany) with a ×20 objective (0.8 NA) or a ×60 oil immersion objective (1.46 NA).

Alexa Fluor 488 and Alexa Fluor 561 were excited with excitation wavelengths of 488 nm and 561 nm, respectively. Alexa Fluor 647 and RBCs were excited with the same excitation wavelength of 647 nm. Three-dimensional imaging data were collected by obtaining images from the x, y, and z-planes. The resulting images were analysed with Imaris software (Bitplane).

### Measurement of meningeal lymphatic vessel diameter distribution

The diameter of lymphatic vessel changed with position. In order to obtain the diameter distribution, we wrote a program with Matlab, which mainly contained two procedures. The schematic diagram was shown as Fig. S4.

#### Procedure 1

This procedure was used to extract the profile of the lymphatic vessel from the initial image. First, Otsu’s method [57] was utilized to decide the threshold and obtain the binary image (Fig. S4-Step 1). Next, image closing operation was used to connect the broken edges of the image. Then, two Matlab functions, “imfill” and “bwareaopen”, were used to fill in the holes and remove the small connected domains of the image, respectively (Fig. S4-Step 2). Finally, the obtained image was subtracted by itself after morphological corrosion, and the profile curve was then determined (Fig. S4-Step 3).

#### procedure 2

This procedure was used to calculate the diameter distribution of lymphatic vessels. As showed in Fig. S4-Step 4, point A and point B represents any points on the outlines on both sides of the lymphatic vessel, and *l*_1_ and *l*_2_ are the tangent lines at point A and B respectively. If *l*_1_, *l*_2_ and *l*_AB_ follows both *l*_1_⊥*l*_AB_ and *l*_2_⊥*l*_AB_, i.e.

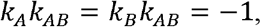

where *k*_A_, *k*_B_ and *k*_AB_ present the slope of line *l*_1_, *l*_2_ and *l*_AB_, respectively. In this case, *l*_AB_ could be taken as the diameter at certain position. However, such point A and B were not always successfully found in all the images. Therefore, for every point A, point B was given by

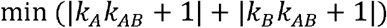

Following above rule, we could obtain a series of |AB| as the lymphatic vascular diameters at every position.

### Fluorescent microscopy monitoring of Evans Blue accumulation in the dcLN

Mice were anesthetized by ketamine/xylazine injection i.p., and then 3 μL of 2% Evans Blue (Sigma-Aldrich) was injected into the right later ventricle (AP: −0.5 mm; DV: 2.5 mm; ML: −1.06mm). 20 min later, the ventral skin of the neck was cut open and dcLNs were exposed. Afterward, a stereo fluorescence microscope (Axio Zoom. V16, Zeiss, Jena, Germany) working at 10× magnification was used for imaging of the dcLNs during the next 1h for each mouse. The mice were treated with PS at same time. Since the Evans Blue could be cleared by MLVs from the brain and be transferred into dcLNs, the accumulation of Evans Blue in dcLNs could reflect the clearance of Evans Blue from brain. After imaging, the fluorescence intensity of the Evans Blue in dcLNs was measured using FIJI software.

### Optical coherent tomography (OCT) monitoring of gold nanorods accumulation in the dcLN

Gold nanorods (GNRs) coated with thiolated polyethylene glycol (0.2 μL, the average diameter, and length at 16±3 nm and 92±17 nm) were injected in the right lateral ventricle (AP: −0.5 mm; DV: 2.5 mm; ML: 1.06 mm). Afterward, OCT imaging of the dcLNs was performed during the next 1h for each mouse.

In this study, a commercial spectral domain OCT Thorlabs GANYMEDE (central wavelength 930 nm, spectral band 150 nm) was used. The LSM02 objective was used to provide a lateral resolution of about 13 microns within the depth of the field. A-scan rate of the OCT system was set to 30 kHz. Each B-scan consists of 2048 A-scans to ensure appropriate spatial sampling.

Since lymph is optically transparent in a broad range of wavelengths, “empty” cavities exist in the resulting OCT image of the lymphatic node with a background signal-to-noise ratio inside. In order to visualize the dynamic accumulation of lymph within these cavities, suspensions of GNRs were used as a contrast agent and the OCT signal intensity is proportional to the GNRs concentration. By tracking the OCT signal temporal intensity changes inside a node’s cavity, we could confirm the clearance pathways and calculate its relative speed. The OCT recordings were performed under anesthesia with ketamine (100 mg/kg, i.p.) and xylazine (10 mg/kg, i.p.).

The GNRs content in the brain tissue and in the dcLNs was evaluated by atomic absorption spectroscopy on a spectrophotometer (Thermo Scientific Inc., Waltham, Massachusetts, USA). The atomic absorption spectroscopy of GNRs was performed 20-40-60 min after the start of OCT monitoring in the brain and in the dcLNs obtained from the same mice, which were used for OCT-GNRs measurements.

### Intracranial pressure monitoring

The ICP monitoring was performed as described previously [58]. For this purpose, the medial part of the left temporal muscle was detached from the skull bone, a small hole was drilled into the temporal bone, and an ICP probe (TSD280) was introduced between the bone and brain into the epidural space and fixed with dental cement. ICP was continuously recorded before, during, and 60 minutes after the IVH on a laptop computer using a micro pressure measurement system (MPMS200), preamplifier (MP150), and software (Biopac, Goleta, CA).

### Tests for evaluation of emotional abnormalities

The post-hemorrhage depression is the most common emotional sequelae of brain hemorrhages, and it’s independently associated with increased morbidity and mortality [59, 60]. The tail suspension, and forced swim test are effective for evaluating emotional abnormality in different models of intracranial hemorrhages [61, 62]. Therefore, these two tests were used for the analysis of early recovery after IVH. The protocol for the tail suspension test was described previously [63, 64]. Briefly, animals were suspended by their tails at the edge of a shelf 55 cm above a desk. Sticky tape (17 cm long) was used to fix the tail (approximately 1 cm from the tip) to the shelf. The recording of mouse mobility and immobility (lack of escape-related behavior when mice hung passively and completely motionless) was made during 360 sec. The protocol of the forced swim test was published in detail in Ref. [65, 66]. The cylindrical tanks (20 cm high, 22 cm in diameter) with water at 24±1°C (10 cm) was used for this test. Each mouse was placed individually in water. The swimming of mice was recorded during 360 sec. The immobility time (when mouse remained floating motionless, making only small movements to keep its head above the water) was calculated during the last 4 min from the 240 s of test time [63].

### Statistical analysis

The results are presented as mean ± standard error of the mean (SEM). Differences from the initial level in the same group were evaluated by the Wilcoxon test. Intergroup differences were evaluated using the Mann-Whitney test and two-way ANOVA (post hoc analysis with Duncan’s rank test). The significance level was set at p < 0.05 for all analyses.

## Acknowledgements

D. Li and S. Liu contributed equally to this work. This work was supported by National Natural Science Foundation of China (NSFC) (Grant Nos. 61860206009, 81870934, 81961138015), the Science Fund for Creative Research Group of China (Grant No. 61721092), the National Key Research and Development Program of China (2017YFA0700501), China Postdoctoral Science Foundation funded project (No. BX20190131, 2019M662633); Funding for Postdoctoral Innovation Research Post in Hubei Province and the Innovation Fund of WNLO; RF Governmental Grant No. 075-15-2019-1885; Russian Science Foundation No. 20-15-00090; Russian Foundation of Basic Research No. 20-015-00308a and No. 19-515-55016 China a. The authors also thank to the Optical Bioimaging Core Facility of WNLO-HUST for support in data acquisition.

## Author contributions

D. Li was involved in the conceptualization, experiment setup, investigation, statistical analysis, writing and editing. S. Liu was involved in the conceptualization, experiment setup and investigation. Z. Liu was involved in statistical analysis. S. Sun was involved in experiment setup and investigation. T. Yu was involved in conceptualization and writing. D. Bragin performed measuring of ICP in mice. N. Navolokin the prepared the human data. J. Kurths was involved in writing and editing. O. Semyachkina-Glushkovskaya and D. Zhu were involved in writing, editing, conceptualization and project management.

## Competing interests

The authors declare no competing interests.

## Notes

### Competing Interest Statement

The authors have declared no competing interest.

## References

1. S. J. An, T. J. Kim, and B. W. Yoon, “Epidemiology, Risk Factors, and Clinical Features of Intracerebral Hemorrhage: An Update,” J Stroke 19(1), 3–10 (2017).

2. P. Daverat, J. P. Castel, J. F. Dartigues, and J. M. Orgogozo, “Death and functional outcome after spontaneous intracerebral hemorrhage. A prospective study of 166 cases using multivariate analysis,” Stroke 22(1), 1–6 (1991).

3. G. Mohr, G. Ferguson, M. Khan et al., “Intraventricular hemorrhage from ruptured aneurysm. Retrospective analysis of 91 cases,” J Neurosurg 58(4), 482–487 (1983).

4. H. E. Hinson, D. F. Hanley, and W. C. Ziai, “Management of intraventricular hemorrhage,” Curr Neurol Neurosci Rep 10(2), 73–82 (2010).

5. W. M. Coplin, F. C. Vinas, J. M. Agris et al., “A cohort study of the safety and feasibility of intraventricular urokinase for nonaneurysmal spontaneous intraventricular hemorrhage,” Stroke 29(8), 1573–1579 (1998).

6. T. Steiner, M. N. Diringer, D. Schneider et al., “Dynamics of intraventricular hemorrhage in patients with spontaneous intracerebral hemorrhage: risk factors, clinical impact, and effect of hemostatic therapy with recombinant activated factor VII,” Neurosurgery 59(4), 767–773; discussion 773-764 (2006).

7. H. Hallevi, K. C. Albright, J. Aronowski et al., “Intraventricular hemorrhage: Anatomic relationships and clinical implications,” Neurology 70(11), 848–852 (2008).

8. K. B. Mahaney, C. Buddhala, M. Paturu et al., “Intraventricular Hemorrhage Clearance in Human Neonatal Cerebrospinal Fluid: Associations With Hydrocephalus,” Stroke 51(6), 1712–1719 (2020).

9. C. Gao, H. Du, Y. Hua et al., “Role of red blood cell lysis and iron in hydrocephalus after intraventricular hemorrhage,” J Cereb Blood Flow Metab 34(6), 1070–1075 (2014).

10. T. S. van Solinge, I. S. Muskens, V. K. Kavouridis et al., “Fibrinolytics and Intraventricular Hemorrhage: A Systematic Review and Meta-analysis,” Neurocrit Care 32(1), 262–271 (2020).

11. H. H. Engelhard, C. O. Andrews, K. V. Slavin, and F. T. Charbel, “Current management of intraventricular hemorrhage,” Surg Neurol 60(1), 15–22 (2003).

12. W. J. Simmonds, “The absorption of blood from the cerebrospinal fluid in animals,” Aust J Exp Biol Med Sci 30(3), 261–270 (1952).

13. M. Oehmichen, H. Gruninger, H. Wietholter, and M. Gencic, “Lymphatic efflux of intracerebrally injected cells,” Acta Neuropathol 45(1), 61–65 (1979).

14. M. Oehmichen, H. Wietholter, H. Gruninger, and M. Gencic, “Destruction of intracerebrally applied red blood cells in cervical lymph nodes. Experimental investigations,” Forensic Sci Int 21(1), 43–57 (1983).

15. M. Caversaccio, O. Peschel, and W. Arnold, “Connections between the cerebrospinal fluid space and the lymphatic system of the head and neck in humans,” in Intracranial and Intralabyrinthine Fluids, pp. PP. 123–128, Springer (1996).

16. W. Dontenwill, “Beitrag zur Genese des Hydrocephalus bzw. der beginnenden Hydranencephalie und zur Frage der Liquorabflußwege,” Frankf Z Pathol 63, (1952).

17. M. Oemichen, “Mononuclear phagocytes in the central nervous system. Origin, mode of distribution, and function of progressive microglia, perivascular cells of intracerebral vessels, free subarachnoidal cells, and epiplexus cells,” Schriftenr Neurol 21, I-X, 1–167 (1978).

18. W. Arnold, R. Ritter, and W. H. Wagner, “Quantitative Studies On The Drainage Of The Cerebrospinal Fluid Into The Lymphatic System,” Acta Oto-Laryngologica 76(1-6), 156–161 (1973).

19. M. Földi, and M. Földi, “Physiologie und Pathophysiologie des Lymphgefäßsystems,” Handbuch der allgemeinen Pathologie 3(6), 239–310 (1972).

20. M. Kozma, Ö. T. Zoltán, and B. Csillik, “Die anatomischen Grundlagen des prälymphatischen Systems im Gehirn,” Cells Tissues Organs 81(3), 409–420 (2008).

21. J. W. Millen, and D. H. M. Woollam, The anatomy of the cerebrospinal fluid, Oxford University Press, Britain (1962).

22. S. Sunderland, “The median nerve anatomical features[” Nerves and Nerve Injuries. Edinburgh: Churchill Livingstone, 754–758 (1972).

23. E. Csanda, “Untersuchungen zur prälymphatischen-lymphatischen zerebro-zervikalen Drainagebahn,” Folia Angiol. 22, 29–31 (1974).

24. E. Csanda, A. Szücs, I. Dobranovics, M. Saal, and A. Oswald, “Hirnschädigungen infolge von Beta-Strahler (Yttrium 90). Weitere Untersuchungen zur Frage der prälymphatisch-lymphatischen Drainagebahn aus der Hirnsubstanz,” Folia Angiol. 1974(22), 172–174 (1974).

25. O. Semyachkina-Glushkovskaya, A. Abdurashitov, A. Dubrovsky et al., “Lymphatic clearance from the blood after subarachnoid hemorrhages,” Proc Spie 10865, (2019).

26. J. M. Chen, L. M. Wang, H. Xu et al., “Meningeal lymphatics clear erythrocytes that arise from subarachnoid hemorrhage,” Nature Communications 11(1), (2020).

27. B. N. Huisa, A. B. Stemer, M. G. Walker et al., “Transcranial laser therapy for acute ischemic stroke: a pooled analysis of NEST-1 and NEST-2,” Int J Stroke 8(5), 315–320 (2013).

28. P. A. Lapchak, and P. D. Boitano, “Transcranial Near-Infrared Laser Therapy for Stroke: How to Recover from Futility in the NEST-3 Clinical Trial,” Acta Neurochir Suppl 121, 7–12 (2016).

29. M. R. Hamblin, “Photobiomodulation for traumatic brain injury and stroke,” J Neurosci Res 96(4), 731–743 (2018).

30. W. Xuan, T. Agrawal, L. Huang, G. K. Gupta, and M. R. Hamblin, “Low-level laser therapy for traumatic brain injury in mice increases brain derived neurotrophic factor (BDNF) and synaptogenesis,” J Biophotonics 8(6), 502–511 (2015).

31. L. D. Morries, P. Cassano, and T. A. Henderson, “Treatments for traumatic brain injury with emphasis on transcranial near-infrared laser phototherapy,” Neuropsychiatr Dis Treat 11, 2159–2175 (2015).

32. M. R. Hamblin, “Low-Level Light Therapy,” (2018).

33. O. Semyachkina-Glushkovskaya, A. Abdurashitov, A. Dubrovsky et al., “Photobiomodulation of lymphatic drainage and clearance: perspective strategy for augmentation of meningeal lymphatic functions,” Biomed Opt Express 11(2), 725–734 (2020).

34. O. Semyachkina-Glushkovskaya, A. Abdurashitov, M. Klimova et al., “Photostimulation of cerebral and peripheral lymphatic functions,” Translational Biophotonics 2(1-2), (2020).

35. A. Dirican, O. Andacoglu, R. Johnson et al., “The short-term effects of low-level laser therapy in the management of breast-cancer-related lymphedema,” Support Care Cancer 19(5), 685–690 (2011).

36. O. Semyachkina-Glushkovskaya, V. Chehonin, E. Borisova et al., “Photodynamic opening of the blood-brain barrier and pathways of brain clearing,” J Biophotonics 11(8), e201700287 (2018).

37. O. Semyachkina-Glushkovskaya, A. Abdurashitov, A. Dubrovsky et al., “Application of optical coherence tomography for in vivo monitoring of the meningeal lymphatic vessels during opening of blood-brain barrier: mechanisms of brain clearing,” J Biomed Opt 22(12), 1–9 (2017).

38. O. Semyachkina-Glushkovskaya, D. Bragin, O. Bragina et al., “Mechanisms of sound-induced opening of the blood-brain barrier,” Adv. Exp. Med. Biol. 1269, (2020).

39. E. Zinchenko, N. Navolokin, A. Shirokov et al., “Pilot study of transcranial photobiomodulation of lymphatic clearance of beta-amyloid from the mouse brain: breakthrough strategies for non-pharmacologic therapy of Alzheimer’s disease,” Biomed Opt Express 10(8), 4003–4017 (2019).

40. C. E. Tedford, S. DeLapp, S. Jacques, and J. Anders, “Quantitative analysis of transcranial and intraparenchymal light penetration in human cadaver brain tissue,” Laser Surg Med 47(4), 312–322 (2015).

41. T. Wang, D. G. Ouzounov, C. Wu et al., “Three-photon imaging of mouse brain structure and function through the intact skull,” Nat Methods 15(10), 789–792 (2018).

42. T. I. Karu, L. V. Pyatibrat, and N. I. Afanasyeva, “Cellular effects of low power laser therapy can be mediated by nitric oxide,” Laser Surg Med 36(4), 307–314 (2005).

43. F. Murad, “Discovery of some of the biological effects of nitric oxide and its role in cell signaling,” Bioscience Rep 24(4-5), 452–474 (2004).

44. J. C. Drapier, H. Hirling, J. Wietzerbin, P. Kaldy, and L. C. Kuhn, “Biosynthesis of nitric oxide activates iron regulatory factor in macrophages,” EMBO J 12(9), 3643–3649 (1993).

45. M. Lepoivre, F. Fieschi, J. Coves, L. Thelander, and M. Fontecave, “Inactivation of ribonucleotide reductase by nitric oxide,” Biochem Biophys Res Commun 179(1), 442–448 (1991).

46. J. C. Drapier, and J. B. Hibbs, Jr., “Aconitases: a class of metalloproteins highly sensitive to nitric oxide synthesis,” Methods Enzymol 269, 26–36 (1996).

47. S. Dimmeler, F. Lottspeich, and B. Brüne, “Nitric oxide causes ADP-ribosylation and inhibition of glyceraldehyde-3-phosphate dehydrogenase,” Journal of Biological Chemistry 267(24), 16771–16774 (1992).

48. J. S. Stamler, D. I. Simon, J. A. Osborne et al., “S-nitrosylation of proteins with nitric oxide: synthesis and characterization of biologically active compounds,” Proc Natl Acad Sci U S A 89(1), 444–448 (1992).

49. Prineas, and J., “Multiple sclerosis: presence of lymphatic capillaries and lymphoid tissue in the brain and spinal cord,” ence 203(4385), 1123–1125 (1979).

50. P. Ballabh, “Intraventricular Hemorrhage in Premature Infants: Mechanism of Disease,” Pediatr Res 67(1), 1–8 (2010).

51. D. Wilson-Costello, H. Friedman, N. Minich, A. A. Fanaroff, and M. Hack, “Improved survival rates with increased neurodevelopmental disability for extremely low birth weight infants in the 1990s,” Pediatrics 115(4), 997–1003 (2005).

52. O. V. Semyachkina-Glushkovskaya, J. Kurths, A. N. Pavlov et al., “Silent Vascular Catastrophes in the Brain in Term Newborns: Strategies for Optical Imaging,” Ieee J Sel Top Quant 22(3), (2016).

53. O. Semyachkina-Glushkovskaya, D. Postnov, T. Penzel, and J. Kurths, “Sleep as a Novel Biomarker and a Promising Therapeutic Target for Cerebral Small Vessel Disease: A Review Focusing on Alzheimer’s Disease and the Blood-Brain Barrier,” Int. J. Mol. Sci. 21(17), 6293 (2020).

54. F. A. Fischbach, D. W. Gregory, P. M. Harrison, T. G. Hoy, and J. M. Williams, “On the structure of hemosiderin and its relationship to ferritin,” J Ultrastruct Res 37(5), 495–503 (1971).

55. B. Akyuz, A. V. Polat, M. Ozturk et al., “Contribution of 3-T Susceptibility-Weighted MRI to Detection of Intraarticular Hemosiderin Accumulation in Patients With Hemophilia,” Am J Roentgenol 210(5), 1141–1147 (2018).

56. A. Louveau, I. Smirnov, T. J. Keyes et al., “Structural and functional features of central nervous system lymphatic vessels,” Nature 523(7560), 337–341 (2015).

57. N. Otsu, “A threshold selection method from gray-level histograms,” IEEE transactions on systems, man, and cybernetics 9(1), 62–66 (1979).

58. D. E. Bragin, R. C. Bush, W. S. Muller, and E. M. Nemoto, “High intracranial pressure effects on cerebral cortical microvascular flow in rats,” J Neurotrauma 28(5), 775–785 (2011).

59. S. Stern-Nezer, I. Eyngorn, M. Mlynash et al., “Depression one year after hemorrhagic stroke is associated with late worsening of outcomes,” Neurorehabilitation 41(1), 179–187 (2017).

60. B. A. Francis, J. Beaumont, M. B. Maas et al., “Depressive symptom prevalence after intracerebral hemorrhage: a multi-center study,” J Patient Rep Outcomes 2(1), 55 (2018).

61. W. Zhu, Y. F. Gao, J. R. Wan et al., “Changes in motor function, cognition, and emotion-related behavior after right hemispheric intracerebral hemorrhage in various brain regions of mouse,” Brain Behav Immun 69, 568–581 (2018).

62. W. Zhu, Y. F. Gao, C. F. Chang et al., “Mouse Models of Intracerebral Hemorrhage in Ventricle, Cortex, and Hippocampus by Injections of Autologous Blood or Collagenase,” Plos One 9(5), (2014).

63. F. Bai, X. Li, M. Clay, T. Lindstrom, and P. Skolnick, “Intra-and interstrain differences in models of “behavioral despair”,” Pharmacol Biochem Be 70(2-3), 187–192 (2001).

64. A. Can, D. T. Dao, C. E. Terrillion et al., “The Tail Suspension Test,” Jove-J Vis Exp (59), (2012).

65. A. Can, D. T. Dao, M. Arad et al., “The Mouse Forced Swim Test,” Jove-J Vis Exp (59), (2012).

66. C. E. Renard, E. Dailly, B. A. Nic Dhonnchadha, M. Hascoet, and M. Bourin, “Is dopamine a limiting factor of the antidepressant-like effect in the mouse forced swimming test?,” Prog Neuro-Psychoph 28(8), 1255–1259 (2004).

