## Supplemental Figure 1-4 for "Photostimulation of lymphatic clearance of red blood cells from the mouse brain after intraventricular hemorrhage"

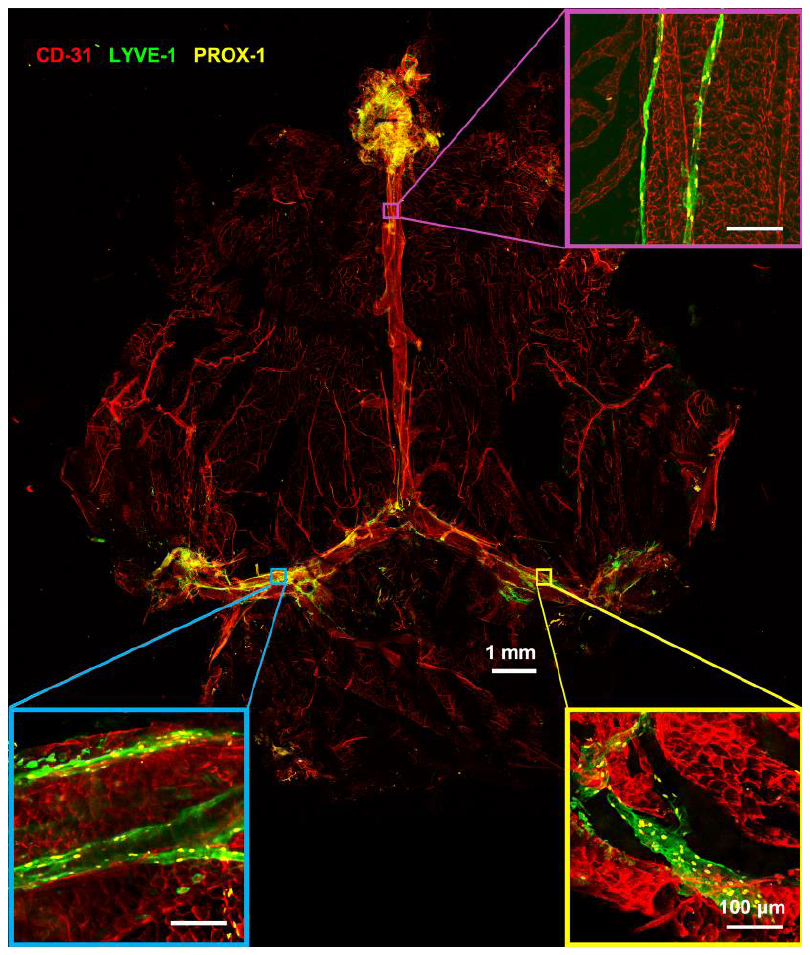

**Fig. S1 Immunofluorescence microscopy of meninge of mice.**

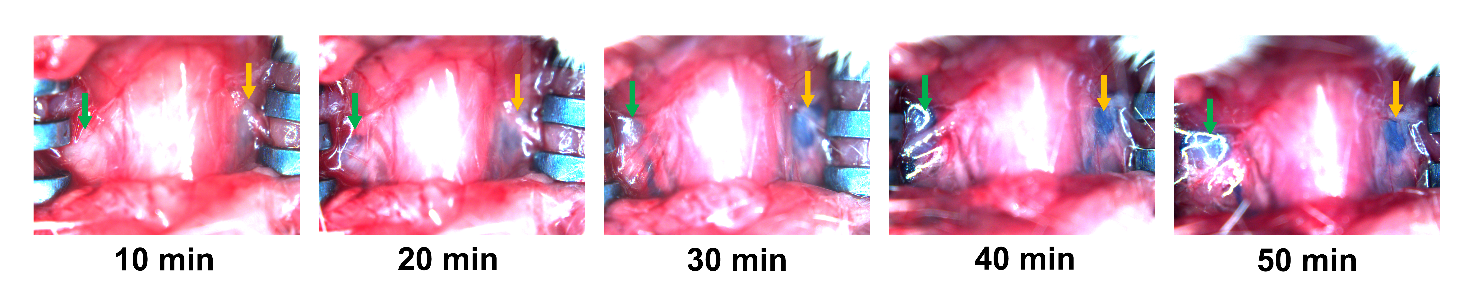

**Fig. S2 Typical photographs of dcLNs post injection of Evans Blue (5%, 5 μL).**

The green and blue arrows represent right and left dcLN, respectively.

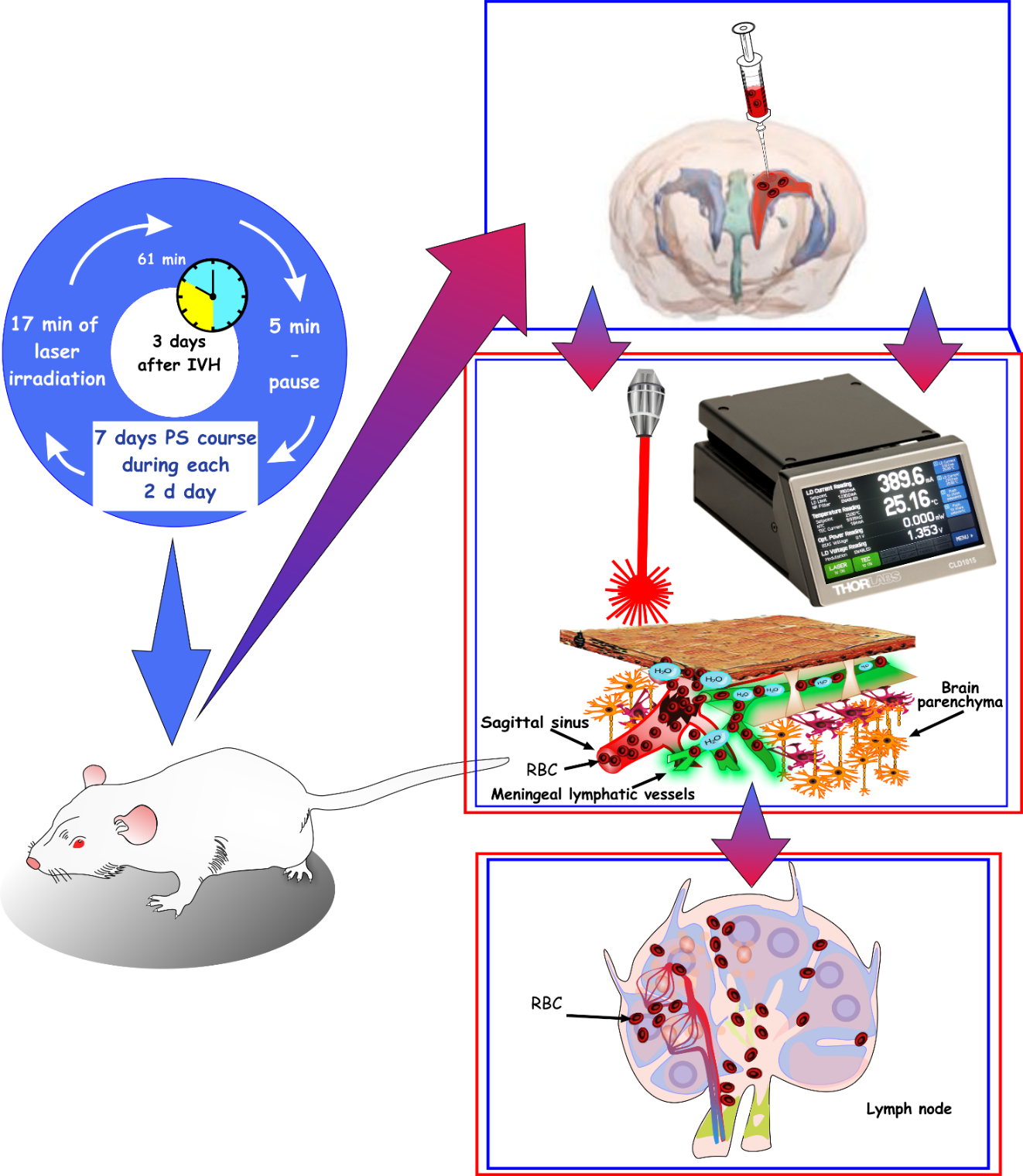

**Fig. S3 Schematic illustration of PS stimulation of RBCs clearance from the mouse brain.**

3 days after a surgical procedure of arterial blood injection into the right lateral ventricle, transcranial PS (1267 nm) of the parietal cortex area in the region of the Sagittal sinus was performed during 7 days each second day under inhalation anesthesia (1% isoflurane at 1 L⁄ min N_2_O⁄O_2_ − 70:30) using the sequence of:17 min – irradiation, 5 min – pause, 61 min in total. The PS course stimulated the RBCs clearance from the brain via MLVs into the dcLNs.

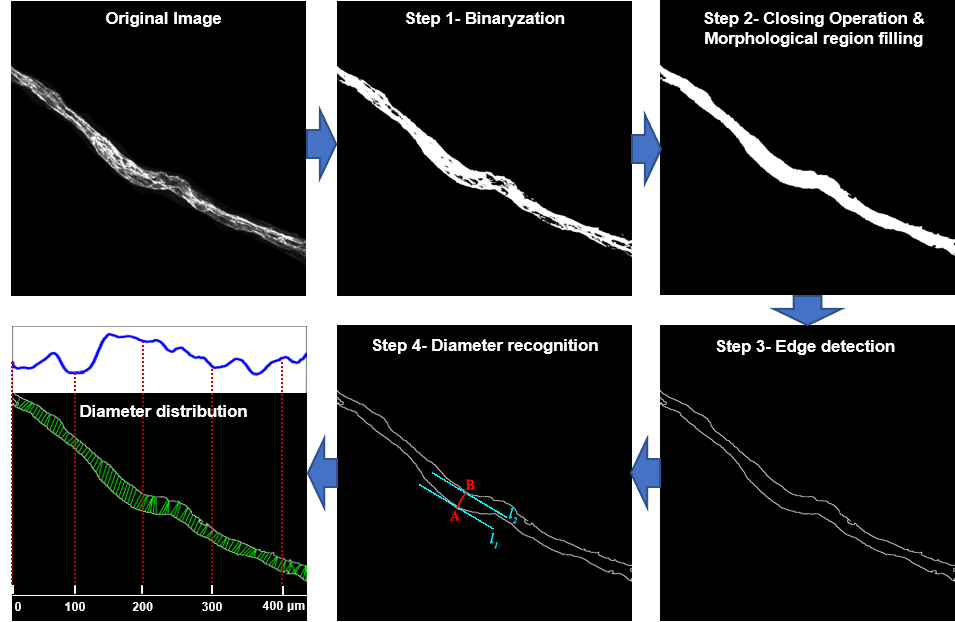

**Fig. S4 Schematic illustration of MLV diameter distribution calculation.**
